# Expression-Based Inference of Cancer Metabolic Flux Differences

**DOI:** 10.1101/2020.01.08.899047

**Authors:** Yiping Wang, Zhenglong Gu

## Abstract

Cancer cells display numerous differences in metabolic regulation and flux distribution from noncancerous cells, which are necessary to support increased cancer cell growth. However, current experimental methods cannot accurately measure such metabolic flux differences genome-wide. To address this short-coming, we apply FALCON, a computational algorithm for inferring metabolic fluxes from gene expression data, to analyze data from The Cancer Genome Atlas (TCGA). We found several major differences between tumor and control tissue metabolism. Cancer tissues have a considerably stronger correlation between RNA-seq expression and inferred metabolic flux, which may indicate a more streamlined and efficient use of metabolism. Cancer metabolic fluxes generally have high correlation with their normal control counterparts in the same tissue, but surprisingly, there are several cases where tumor samples in one tissue have even higher correlation with control samples in another tissue. Finally, we found several pathways that frequently have divergent flux between tumor and control samples. Among these are several previously implicated in tumorigenesis, including sphingolipid metabolism, methionine and cysteine synthesis, and bile acid transformations. Together, these findings show how cancer metabolism differs from normal tissues and may be targeted in order to control cancer progression.

## 2 Introduction

Alterations in metabolism have recently been identified as one of the hallmarks of cancer [1]. This recognition comes in the wake of numerous studies showing that cancer cell metabolism is broadly dysregulated [2]. Indeed, many common driver mutations have been shown to cause coordinated changes in cancer signaling and metabolic networks, reprogramming their metabolism in order to support the demands of continued proliferation [2]. In normal cells, proliferation takes place when growth factors bind to receptors such as IGFR, EGFR, and PDGFR, leading to upregulation of pro-growth signaling pathways such as PI3K-Akt, mTOR, and c-Myc [3] [4] [5]. Mutations in either these receptors, or components of the downstream signaling pathway proteins, can lead to changes in their activity patterns in cancer cells. In turn, this leads to changes in metabolic regulation, including but not limited to changed expression and degradation rates of metabolic enzymes and transporters, different patterns of phosphorylation and acetylation, and altered concentrations of allosteric regulators such as fructose-2,6-bisphosphate [2].

These regulatory changes are believed to cause a rewiring of metabolism in cancer, so that flux through metabolic pathways is reorganized in order to support increased cell proliferation [6]. The oldest and most well-studied of these pathway changes is the Warburg effect, first observed by Otto Warburg, in which cancer cells uptake glucose and use glycolysis to ferment it, and excrete the resulting lactate at very high rates, even if they have access to normal oxygen levels [7]. Despite glycolytic fermentation generating much less ATP than aerobic respiration, the Warburg effect does allow more glucose to be used for production of biosynthetic precursors, such as nucleic and amino acids [6]. These compounds are also needed for rapid cancer cell growth, and often are the limiting factor in such growth instead of ATP [6]. In addition, several other pathways, including glutaminolysis [8] and one-carbon metabolism [9], have been intensively studied in recent years, and have similarly been shown to contribute significantly to tumorigenesis.

Yet there are still many details of cancer metabolism that remain to be explored. Most importantly, there are many additional metabolic pathways that have yet to be experimentally studied for their impact on tumorigenesis. Partly, this is because the most common experimental method for studying experimental is 13C flux analysis [10]. This method uses nutrients, such as glucose and glutamine, which have been labeled at specific positions with 13C atoms in place of 12C. By feeding such nutrients to cells in cell culture, the labeled nutrients are broken down and 13C atoms dispersed so that they label other metabolic compounds. These labeling patterns can then be measured through mass spectrometry, and metabolic flux can be computationally inferred from them. But although 13C flux analysis can give very clear results for some pathways, it has not yet been scaled up to a level that can measure metabolism across an entire cell [11].

We attempt to address this problem, by using a computational method previously developed in our lab called FALCON [12]. FALCON is part of a general class of computational methods called constraint-based modeling (CBM) [13], which model the interactions in a metabolic network as a matrix, and model constraints on metabolic flux, such as the availability of nutrients to a cell, using a series of linear inequalities. Using these concepts, FALCON then takes as input metabolic gene expression measurements from cells, maps the expression onto the metabolic network, and maximizes the linear correlation between expression and predicted metabolic flux. FALCON thus relies on the assumption that high metabolic enzyme expression is correlated with high flux, and vice versa, as described further in Methods.

**Table 1:** TCGA datasets used in this study.

| Dataset | Number of Matched Tumor and Control Samples |
| --- | --- |
| BLCA | 19 |
| BRCA | 113 |
| CHOL | 17 |
| COAD | 41 |
| ESCA | 11 |
| HNSC | 44 |
| KICH | 49 |
| KIRC | 72 |
| KIRP | 32 |
| LIHC | 50 |
| LUAD | 59 |
| LUSC | 49 |
| PRAD | 52 |
| READ | 11 |
| STAD | 32 |
| THCA | 58 |
| UCEC | 23 |

We thus applied FALCON to study metabolic flux in cancer cells vs. normal controls, using data from The Cancer Genome Atlas (TCGA) [14]. Our goal was to determine which metabolic pathways were most frequently altered in cancer cells, and the pattern of such alterations across the metabolic network.

## 3 Methods

We downloaded manifest files for FPKM files of mRNA expression on 17 major tissues from the GDC data portal at https://portal.gdc.cancer.gov. We chose tissues which had a minimum of 10 matched tumor and normal control samples, in order to carry out pairwise statistical tests later on. The 17 tissues are listed in 1, along with the corresponding number of samples for both cancer tissues and normal controls. For our analysis, we used only samples that were marked as either Primary Tumor Sample or Solid Tumor Control in our data. This excludes any samples that were marked as Metastatic.

We then mapped expression from our downloaded FPKM files onto the latest version of the human metabolic reconstruction, Recon3.02 [15]. This reconstruction models 7440 metabolic reactions that have been found in human cells. We also applied constraints to Recon2 in order to model limitations on the nutrients available to human cells. We were unable to find a source for the composition of the extracellular medium that would be located in each tissue. Therefore, we developed our own list of constraints that is intended to model the nutrients that would be available to cells grown in cell culture medium. For 43 nutrients which are listed in 2, chiefly amino acids, vitamins and ions, we allowed unlimited uptake or excretion into our model. For all remaining metabolites, we did not allow uptake from the extracellular medium, as we were uncertain if they were present at significant levels, but allowed possible excretion if internal reactions in our model produced them.

**Table 2:** Recon2 metabolites that were allowed to be uptaken by the model in this study.

| Metabolite Name | Metabolite Abbreviation in Recon2 |
| --- | --- |
| glycine | gly\|e\| |
| L-argininium | arg_\|L\|e\| |
| L-aspartate | asp_\|L\|e\| |
| L-asparagine | asn_\|L\|e\| |
| L-cysteine | cys_\|L\|e\| |
| L-glutamate | glu_\|L\|e\| |
| L-glutamine | gln_\|L\|e\| |
| L-histidine | his_\|L\|e\| |
| trans-4-hydroxy-L-proline | 4hpro_\|LT\|e\| |
| L-isoleucine | ile_\|L\|e\| |
| L-leucine | leu_\|L\|e\| |
| L-lysine | lys_\|L\|e\| |
| L-methionine | met_\|L\|e\| |
| L-phenylalanine | phe_\|L\|e\| |
| L-proline | pro_\|L\|e\| |
| L-serine | ser_\|L\|e\| |
| L-threonine | thr_\|L\|e\| |
| L-tryptophan | trp_\|L\|e\| |
| L-tyrosine | tyr_\|L\|e\| |
| L-valine | val_\|L\|e\| |
| biotin | btn\|e\| |
| choline | chol\|e\| |
| (R)-pantothenate | pnto_\|R\|e\| |
| folate | fol\|e\| |
| nicotinamide | ncam\|e\| |
| benzoate | bz\|e\| |
| pyridoxine | pydxn\|e\| |
| riboflavin | ribflv\|e\| |
| thiamine | thm\|e\| |
| myo-inositol | inost\|e\| |
| calcium(2+) | ca2\|e\| |
| sulfate | so4\|e\| |
| potassium | k\|e\| |
| chloride | cl\|e\| |
| sodium | na1\|e\| |
| bicarbonate | hco3\|e\| |
| hydrogen phosphate | pi\|e\| |
| reduced glutathione | gthrd\|e\| |
| oxygen | o2\|e\| |
| carbon dioxide | co2\|e\| |
| water | h2o\|e\| |
| proton | h\|e\| |

### 3.1 Description of constraints-based modeling and FALCON

Metabolic networks can be considered as bipartite networks, in which each reaction is linked to the metabolites that it consumes and/or produces [13]. CBM methods then model a metabolic network as a stoichiometric matrix *S*, in which each row represents a metabolite, each column a reaction, and at each row-column intersection is a coefficient representing how many molecules of a metabolite are involved in each reaction. Using this matrix, two major physicochemical constraints may be imposed. First, at steady-state, the concentrations of metabolites in a cell are neither rising or falling. Therefore, given a vector of fluxes defined as *v*, this constraint may be written as the equation *S · v* = 0. Second, all metabolic reactions have a maximum rate, given that a cell can produce only a finite amount of enzymes, and enzymes’ efficiency is limited. This constraint may be written as *v* <= *v*_*max*_, where *v*_*max*_ is the maximum rate of a metabolic reaction.

In the case of some reactions, *v*_*max*_ can be inferred based on maximum levels of experimentally measured flux rates. This is especially important for reactions which model the uptake of nutrients such as oxygen or glucose, which are often the limiting factor that constrains cell growth rates. For other reactions, *v*_*max*_ is set to an arbitrary large value, usually + − 1000. This is done so that flux through these reactions will not become the limiting factor in cell growth rates.

Traditionally, metabolic network flux distributions are inferred using flux balance analysis with a biomass objective. A biomass objective represents the proper ratios of amino acids, nucleotides, and so forth necessary to create biomass, and a flux distribution that maximizes it could be said to maximize cell growth rate and therefore fitness. However, biomass optimization is generally not considered the be the most accurate way to model human cell metabolism [16]. Each tissue in humans is known to carry out distinct functions for which its metabolism is tailored, such as ion exchange in kidney cells, neuromodulator production in nerve cells, etc. During the process of tumorigenesis, cancer cells may modify normal tissue functions in order to support increased growth rates. However, mRNA and protein expression analyses of the TCGA have shown that tumor samples still retain considerable similarity to corresponding control tissues [17], and therefore metabolism of cancer tissues is also likely to retain tissue specificity. To capture this specificity more accurately, we therefore applied FALCON, which is a previously developed method in our lab that infers flux based on gene expression, by optimizing the correlation between flux and gene expression. Biomass rate is not optimized under FALCON, but instead, FALCON relies upon the assumption that high gene expression is generally correlated with high metabolic flux, and vice versa. The degree to which this assumption is true varies depending on the specific cell type and pathway. Previous investigations in bacteria have shown at least some cases where it is true, and on this basis, we have used FALCON to analyze human TCGA data as well [18] [19].

## 4 Results

### 4.1 Expression-Flux Correlations in Tumor and Control Samples

For each of the 17 tissues we examined, we first wanted to check the correlation of measured expression values with predicted flux in our FALCON simulations. FALCON is designed to maximize linear correlation of predicted flux values to expression. However, due to constraints imposed by the topology of the metabolic network, we generally find that this correlation is poor. For example, among all control samples taken from breast tissue in the TCGA dataset, we calculated the average expression and predicted flux of all 7440 reactions in the Recon2 model. We then calculated the Spearman’s correlation between average expression and flux, for all reactions with either nonzero average expression or flux. There are 5518 such reactions, with a Spearman’s *R*^2^ of just .0016 (Figure 1a). Similar results hold for breast tumor samples (Figure 1b). However, among breast control samples, 5563 reactions in the model have either zero average expression or flux. These reactions are located at the bottom and left edges of Figure 1a. The presence of these two conflicting groups causes low expression-flux correlation, if they are included in calculating correlation. If they are excluded, then among the 1877 remaining reactions, flux-expression correlation rises to an *R*^2^ of .244. In other words, although flux-expression correlation is globally very low across the metabolic network, there is significant flux-expression correlation among the subset of the network that consistently has nonzero metabolic expression and flux.

**Figure 1:**
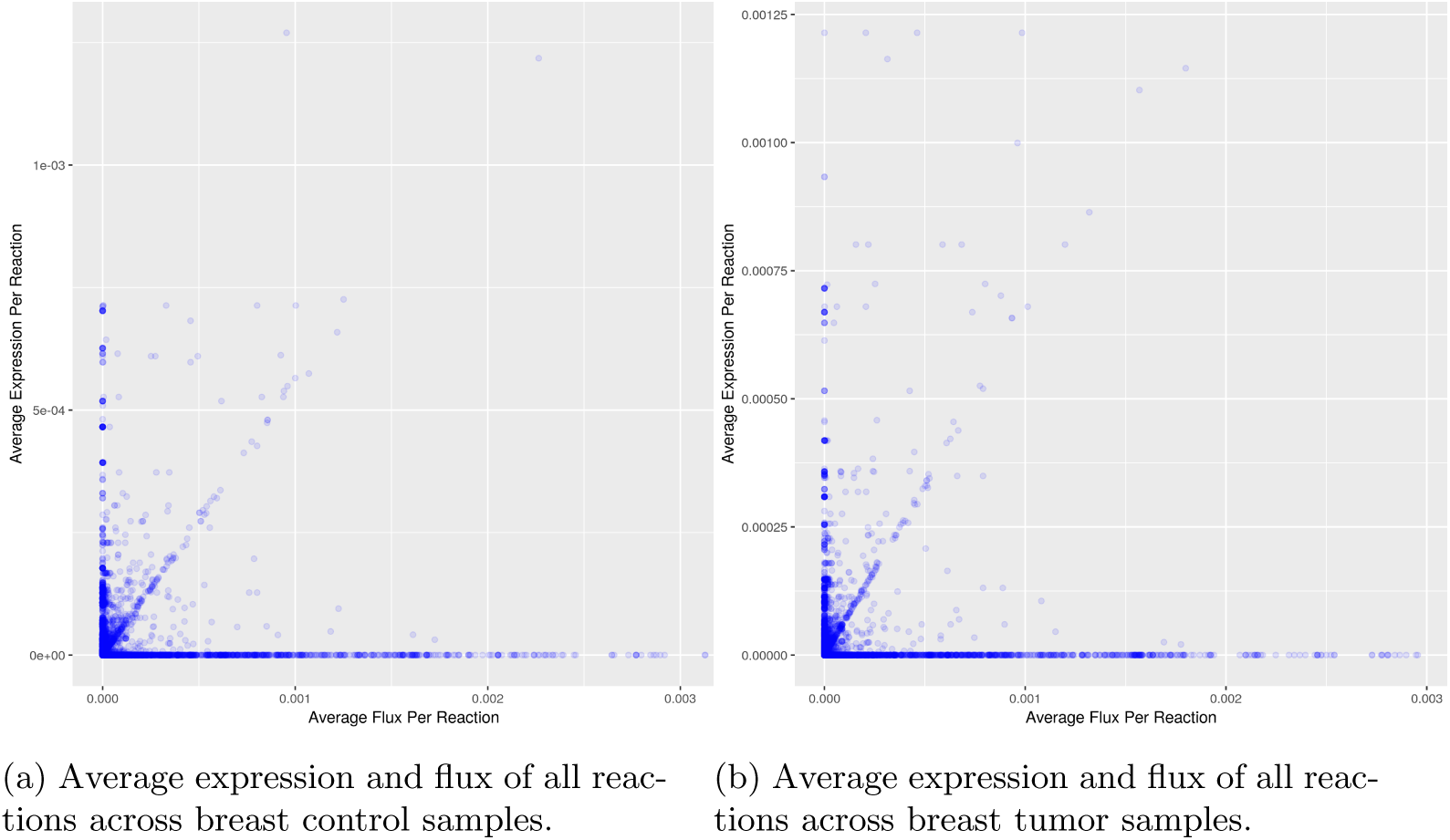
Expression vs. flux correlation for breast samples.

Furthermore, we observed an interesting pattern across all TCGA datasets when we compared flux-expression correlations in tumor samples vs. normal samples (Figure 2). These correlations are significantly stronger in tumor samples than normal samples, across almost all datasets. When considering reactions with nonzero flux OR nonzero expression, the increase is only .0017 on average, although it applies to all datasets besides cholangiocarcinoma. However, for reactions with nonzero flux AND nonzero expression, the average increase is .083, and all datasets show such an increase except for prostate. Furthermore, the total number of reactions with nonzero expression OR nonzero flux is on average 529 reactions greater in control vs. tumor samples. For nonzero expression AND flux, the difference is on average 434 more reactions in control samples.

**Figure 2:**
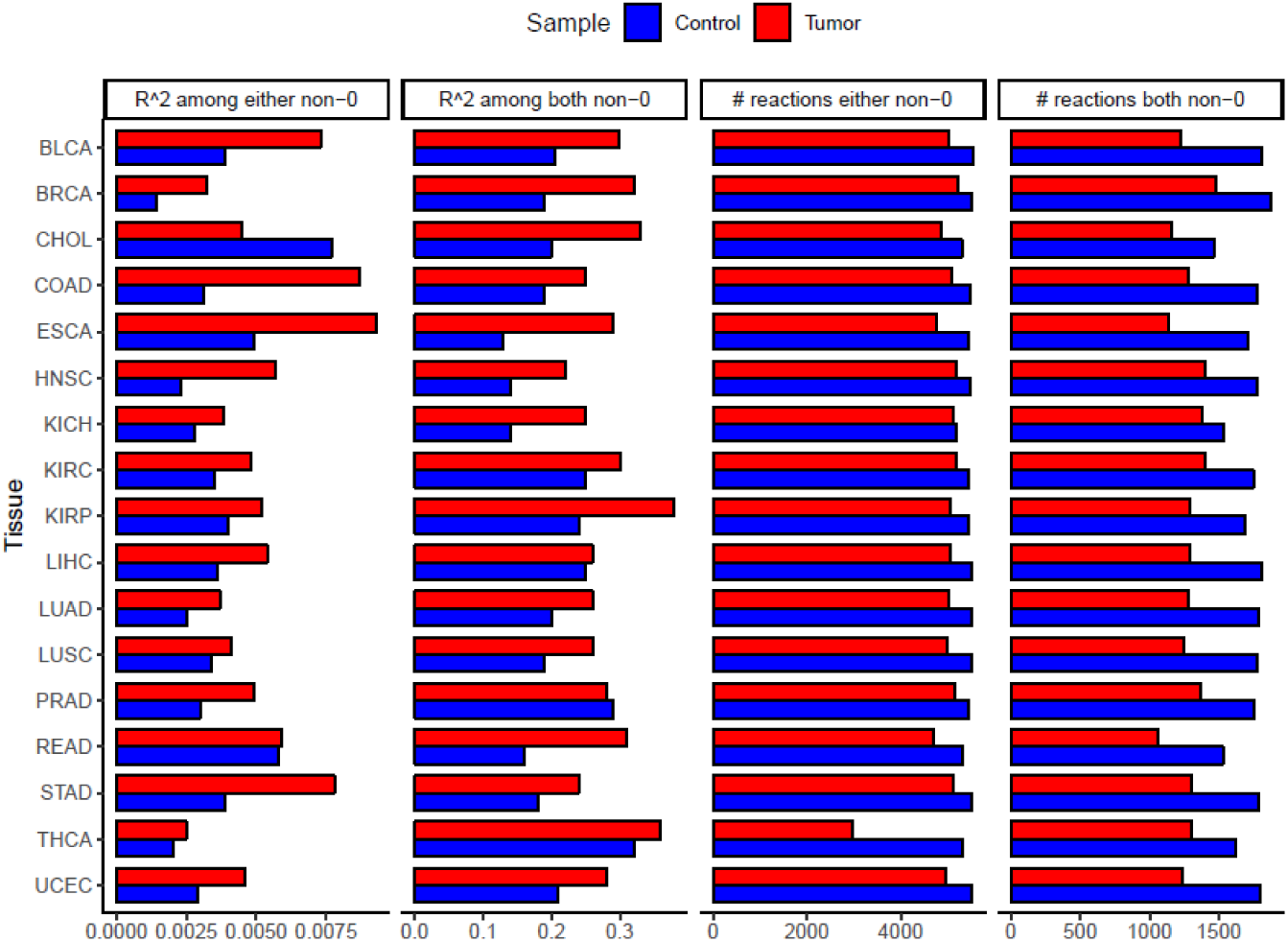
Expression-flux correlations and numbers of nonzero reactions, across 17 TCGA tissues.

Our interpretation of these results is that tumor samples exhibit a more streamlined metabolism, with less post-transcriptional regulation than control samples. Greater expression-flux correlation indicates that regulation after transcription, such as differences in translation rate or post-translational modifications, has less impact in weakening the correlation between expression flux. This may occur because tumor tissues have large numbers of mutations throughout their genome [20], some of which may alter post-transcriptional regulation of metabolism. Furthermore, if fewer reactions have nonzero expression and flux in tumor samples, this indicates that tumor samples have switched off some reactions that are unnecessary for proliferation. This makes sense, considering that normal tissues are specialized to carry out distinct metabolic functions, whereas cancers undergo strong positive evolution for the sole purpose of proliferating quickly, and therefore may converge on a common metabolic phenotype [21].

### 4.2 Flux-Flux and Expression-Expression Correlations in Tumor and Control Samples

We furthermore wanted to examine how similar flux and expression may be in normal and tumor samples of the same tissue. As an example, we plotted the average flux of each reaction in breast control and tumor samples, and calculated the Spearman correlation between the two measurements (Figure 3). We did the same procedure for expression measurements as well. Both analyses showed high correlations of *R*^2^ = .89 for flux and .98 for expression. Furthermore, this pattern applies to all of our examined datasets, with high correlations for both flux-flux and expression-expression comparisons, and the latter being stronger. This implies that although overall metabolic expression differences between tumor and control samples may be small, they result in somewhat greater differences in flux later on. As will be seen, we also find a number of differential flux and expression subsystems in each tissue.

**Figure 3:**
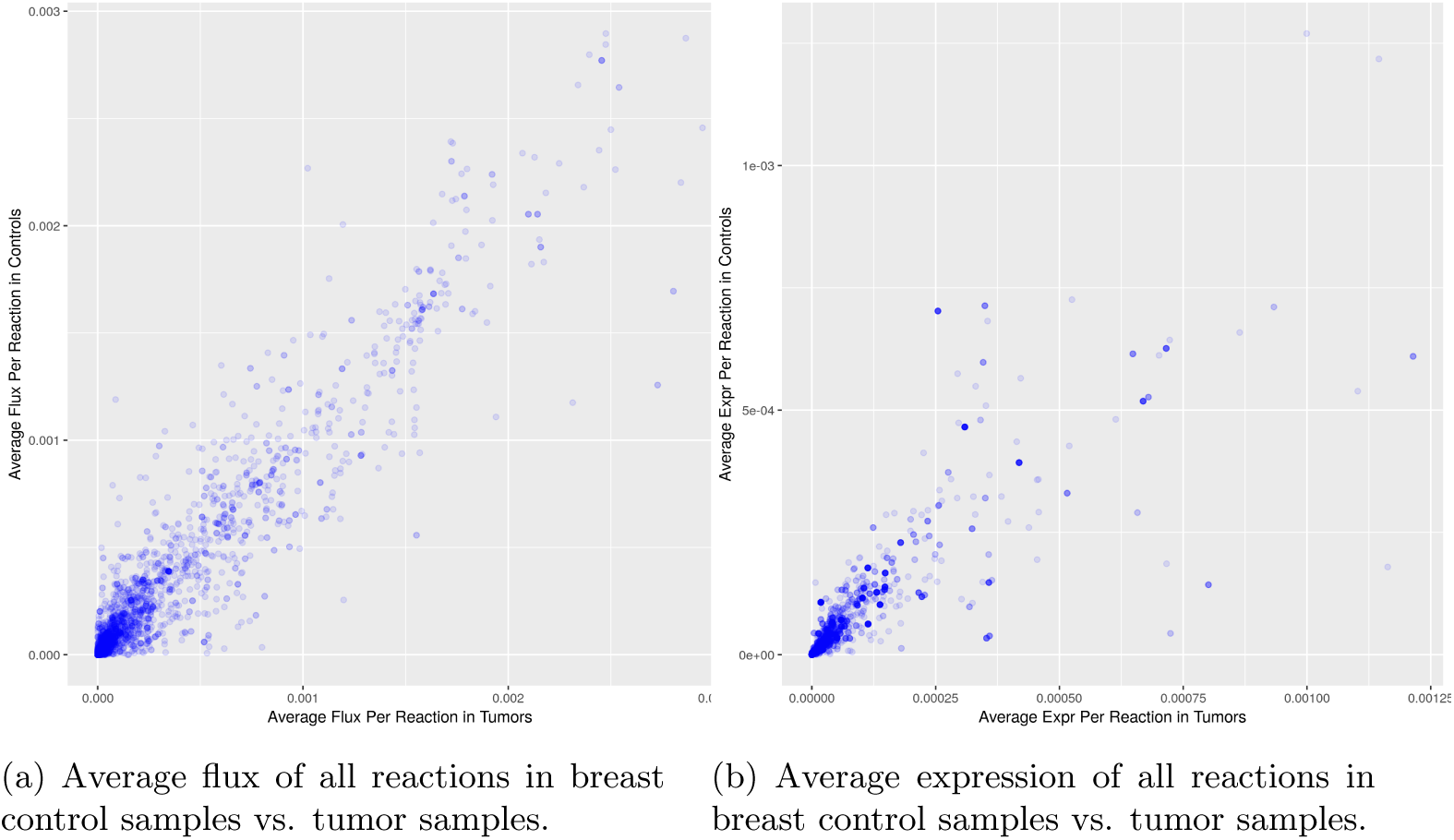
Flux vs. flux and expression vs. expression correlations for breast samples.

We also compared the flux-flux and expression-expression correlations between tumor samples in one tissue, versus control samples in another tissue (Figure 4). To our surprise, we find that in most cases, tumor samples do not show the highest flux or expression correlation with control samples in the same tissue, but instead with control samples in some other tissue. For example, tumor cholangiocarcinoma samples have a flux-flux correlation of .76 with bile duct control samples, but a correlation of .80 with liver control samples in the LIHC dataset. Although the difference is not large, this unexpected result implies that in the course of tumorigenesis, tumor samples alter their metabolic expression and flux, in such a way that it actually results in higher similarity to a different control tissue than the original tissue. Furthermore, there are a few cases where the two tissues are closely related, such as LUSC (lung squamous cell carcinoma) and LUAD (lung adenocarcinoma), but the majority of connections are between unrelated tissues.

**Figure 4:**
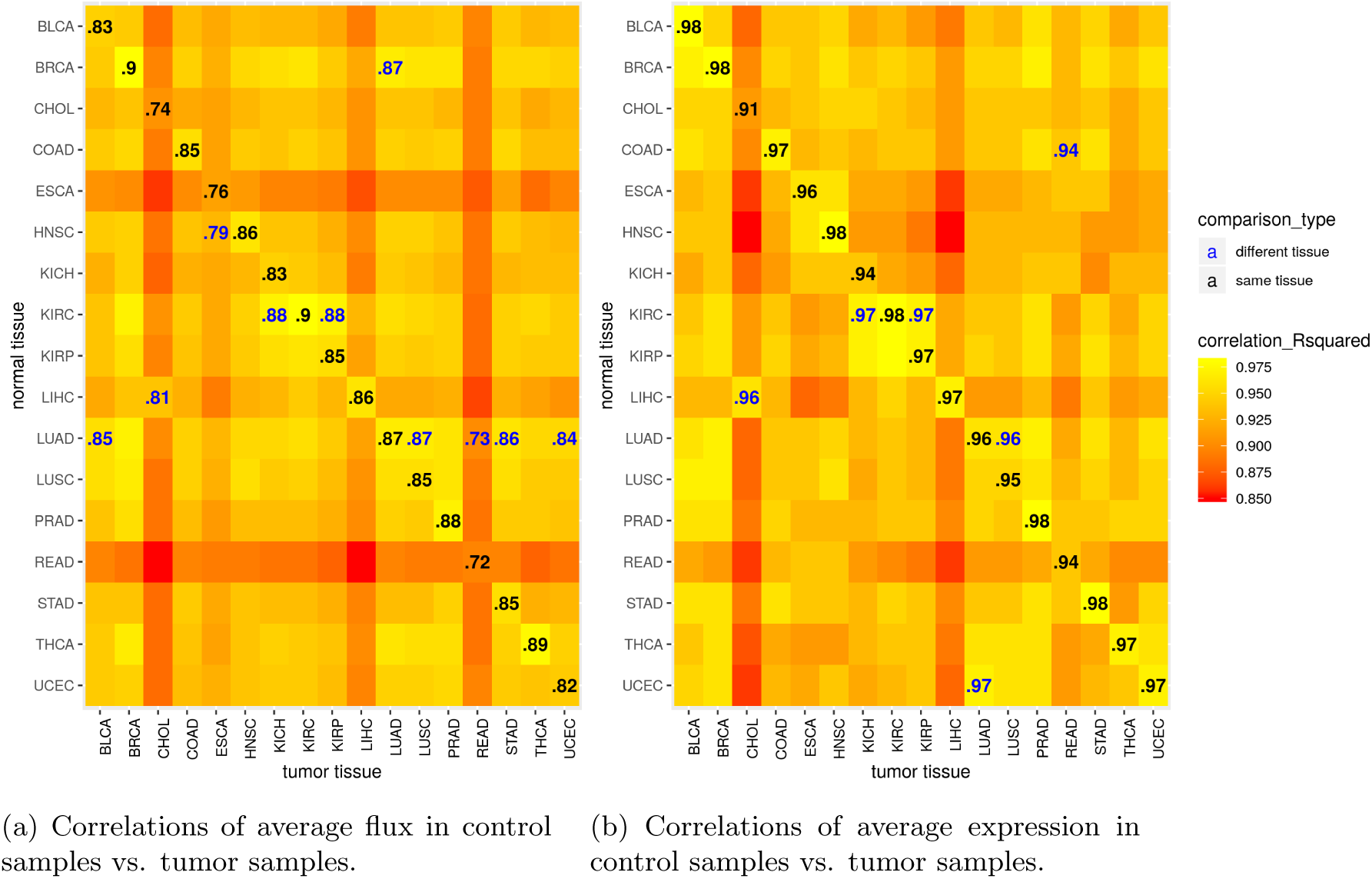
Correlations of tumor flux vs. control flux and tumor expression vs. control expression across all 17 TCGA tissues.

### 4.3 Differential Metabolic Flux and Expression Subsystems

For each of the 17 tissues we examined, we used the Wilcoxon signed-rank test to determine reactions with significant differential expression or flux between tumor samples vs. normal controls. We then determined metabolic subsystems that were enriched in such reactions, by using gene set enrichment analysis (GSEA) [22]. We also repeated this analysis with expression data instead of flux data for each sample, to calculate differential expression for all reactions and pathways in each tissue. For both of these analyses, we considered a subsystem to be enriched in a tisue if its GSEA p-value was less than .05.

Overall, thirty metabolic subsystems were enriched in reactions with higher flux in tumors in at least 1 tissue. Our results highlight several pathways with high differential flux that are already known to be important in tumorigenesis. The most common differential flux subsystem with higher flux in tumors, in eight out of seventeen tissues, is methionine and cysteine metabolism, which involves reactions for biosynthesis of these two closely related amino acids. Both of them are required inputs to folate and one-carbon metabolism, which are further linked to DNA methylation and nucleotide synthesis pathways. A recent paper has shown that methionine restriction has an anti-tumorigenic effect [23], and therefore it is not surprising that higher biosynthetic flux is observed in tumors. Furthermore, the next most common subsystem with higher flux in tumors is tyrosine metabolism, which also involves biosynthesis of a key essential amino acid. Fatty acid synthesis has higher tumor flux, because it is necessary for increased cell membrane size, which expands greatly in area to accompany cell proliferation in tumors. Sphingolipid metabolism also had higher differential flux in several tissues, which may be related to the properties of sphingolipids as key signaling metabolites, with either an pro- or anti-tumorigenic effect in many tumors [24]. Production of the sphingolipid ceramide is associated with increased apoptosis, autophagy, and cell death, while the related sphingolipid sphingosine-1-phosphate activates oncogenic signaling receptors [24]. Finally, bile acid metabolism shows higher flux metabolism in tumors, which is highly interesting given a recent paper on how the gut microbiome may modify primary bile acids that are initially produced by the liver into secondary bile acids. These secondary bile acids have been shown to promote liver tumorigenesis by altering the immune response in liver tissues [25].

Twenty-six subsystems were enriched in reactions with lower tumor flux, in at least one tissue. Among the most common subsystems among all tissues are nucleotide interconversion and the pentose phosphate pathway. Both of these are involved in the supply of nucleotides for DNA synthesis, so it is surprising that they exhibit lower flux in tumors. However, one possibility is that tumors instead obtain most of these compounds by scavenging from the extracellular environment, which has been previously been shown to be a major contributor to cancer metabolism [26]. Similarly, although cholesterol metabolism flux is lower in most tumors compared to controls, this may be explained by higher tumor scavenging of cholesterol. Cholesterol is a major component of cell membranes, and is also important in cancer signaling related either to lipid rafts or mTORC1 [27]. On the other hand, N-glycan synthesis, aminosugar metabolism, and inositol phosphate metabolism all exhibit lower flux, and in this case may be expected. All three subsystems are related to the synthesis of glycoproteins which frequently serve as signaling markers on the cell surface, and the composition of glycoproteins has been shown to be altered in cancers [28]. Finally, starch and sucrose metabolism, fructose and mannose metabolism, and pyrimidine metabolism are reduced as well, further suggesting that cancers may reduce their dependence on these nutrient sources, in favor of others.

Furthermore, when we look at differential expression subsystems in our dataset, we saw first there were considerably fewer such subsystems, with only seventeen with lower tumor expression in at least one tissue, and fourteen with higher. Nevertheless, a few of these overlap with the differential flux subsystems previously described, among them sphingolipid metabolism and bile acid synthesis with higher flux and expression in tumors, and nucleotide interconversion, pyrimidine catabolism, N-glycan synthesis and aminosugar metabolism with lower flux and expression. These results give us added confidence that the subsystems in question are indeed upregulated in cancer metabolism, and play a significant role in tumorigenesis. On the other hand, the remaining differential flux subsystems, which do not have differential expression, indicate that FALCON may be able to independently predict significant flux differences, and thus give additional information on metabolism, outside of expression measurements. But furthermore, there are also several subsystems with strong differential expression, but without corresponding differential flux. Among those with higher tumor expression are tryptophan metabolism, which involves an essential amino acid, and steroid metabolism, which may be important in cancer signaling networks. Folate metabolism has lower tumor expression, though it has been shown to be essential in maintaining tumor nucleotide synthesis and methylation. These subsystems probably do correspond to real cases of differential flux in cancer tissues, but they are not predicted by FALCON. One reason may be that mRNA expression is known to not be perfectly correlated with flux [19], and FALCON may therefore perform better if given access to proteomics data, which is believed to be more relevant for flux prediction.

To further test whether we obtain significant flux and expression differences between tumor and control samples, we also used random forests to extract significant reactions. We used the package randomForests in R, with the parameters ntrees=501, importance=TRUE, proximities=501, to distinguish normal from tumor samples in all 17 tissues. From the out-of-bag error of the trees, we found that classification accuracy was very high in all tissues, using both flux and expression data, with an average of 96% accuracy. However, if we rank all metabolic reactions by their importance in classifications, according to mean decrease in accuracy when the reaction is excluded, we find that there is a very smooth decline in importance. We therefore decided to rank metabolic subsystems based on their average importance. We calculated the average importance of reactions within each subsystem, and used this to rank each subsystem.

Importantly, there is a weak, but significant negative correlation, between the average importance of a subsystem according to random forests, and its p-value in GSEA. Specifically, for each subsystem in Recon2, and each tissue for which control and tumor samples, we obtained both the average importance of its reaction fluxes in random forests, as well as its p-value in GSEA. We computed the correlation of these two measures, as well as for expression measurements. The results are plotted in Figure 6, and show that the Spearman’s *R*^2^ values are .07 for flux correlation and .15 for expression measurements, which although low have corresponding p-values of 3.80e-12 and 9.23e-28.

**Figure 5:**
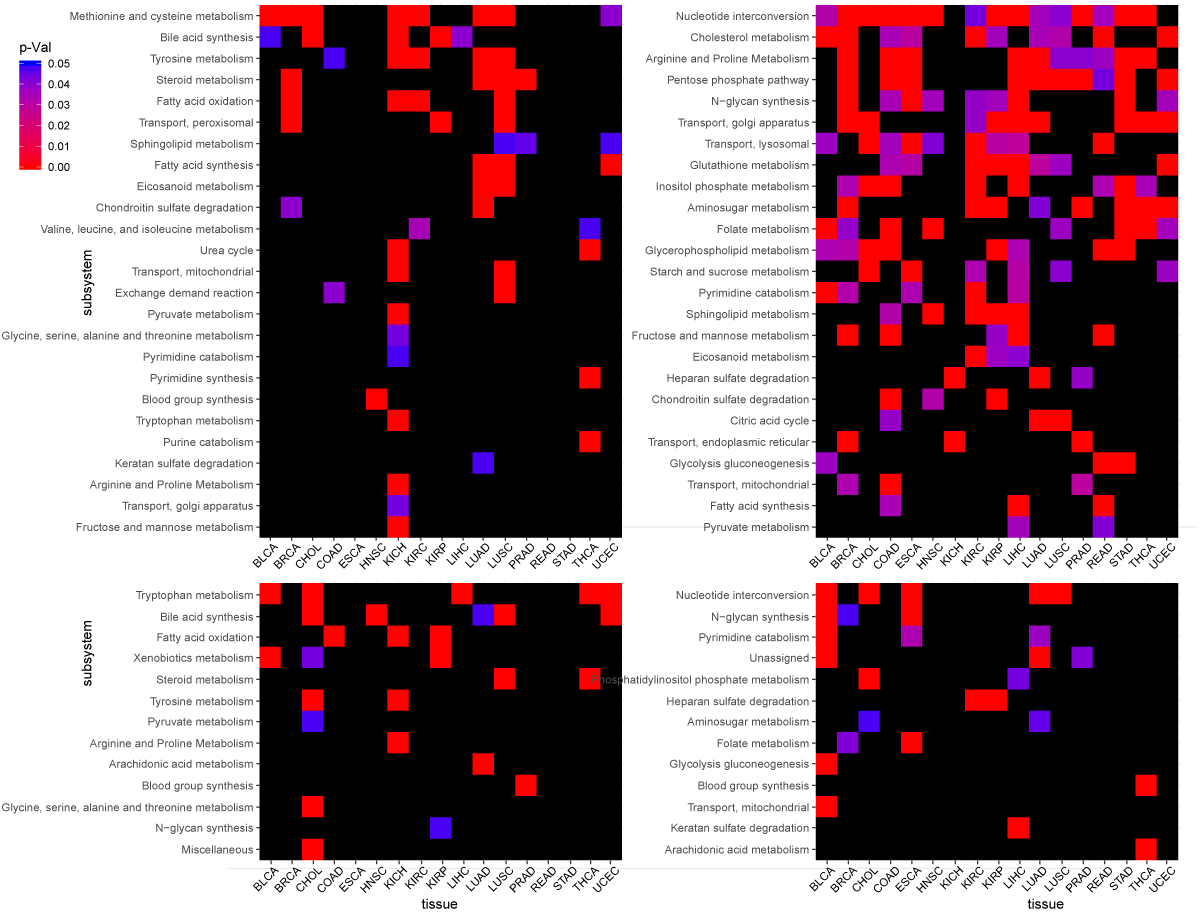
Top-left: GSEA positive differential flux subsystems. Top-right: GSEA negative differential flux subsystems. Bottom-left: GSEA positive differential expression subsystems. Bottom-right: GSEA negative differential expression subsystems.

**Figure 6:**
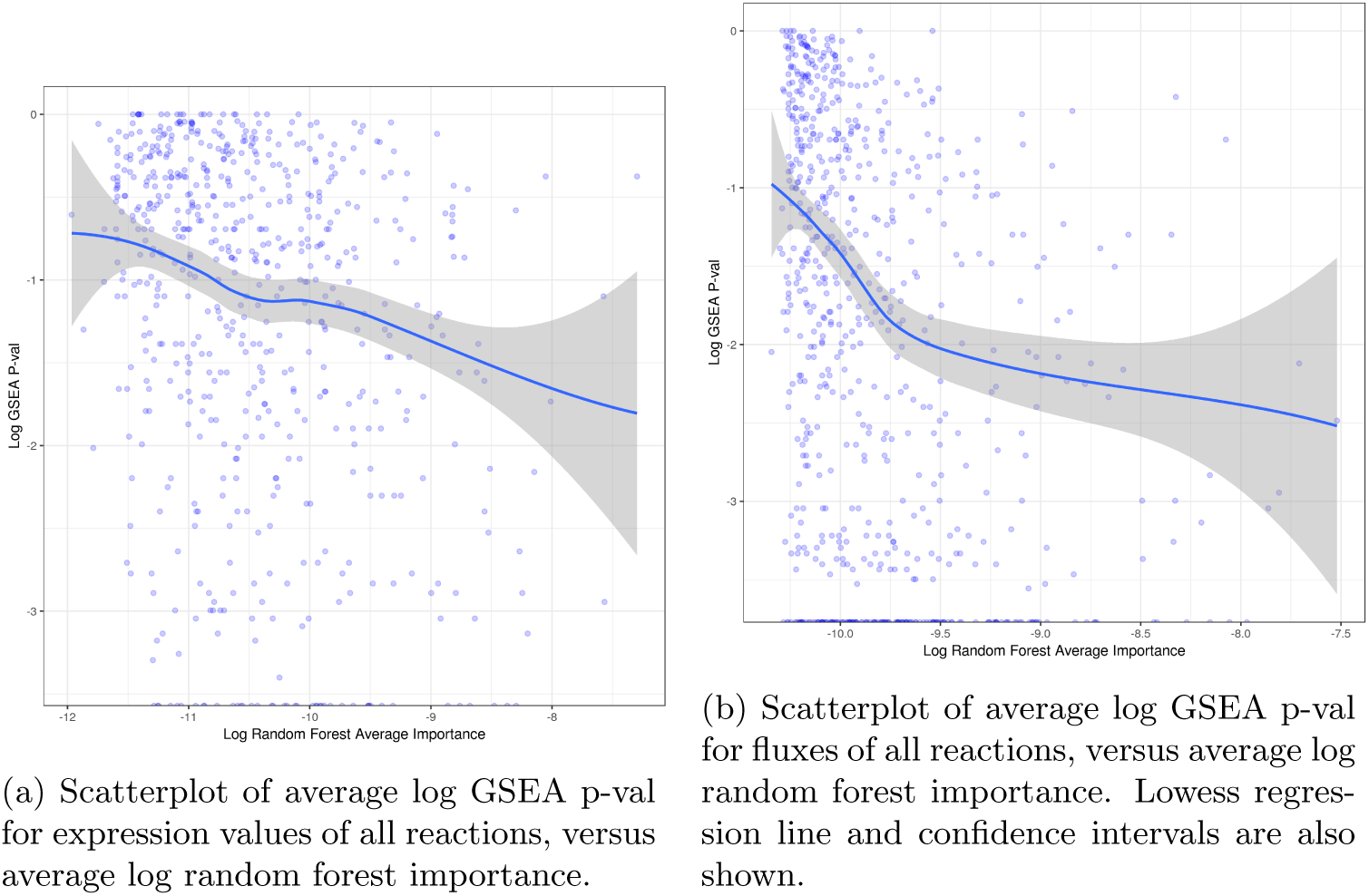
Scatterplots of average log GSEA p-val versus average log random forest importance, for either expression values or fluxes of all reactions.

**Figure 7:**
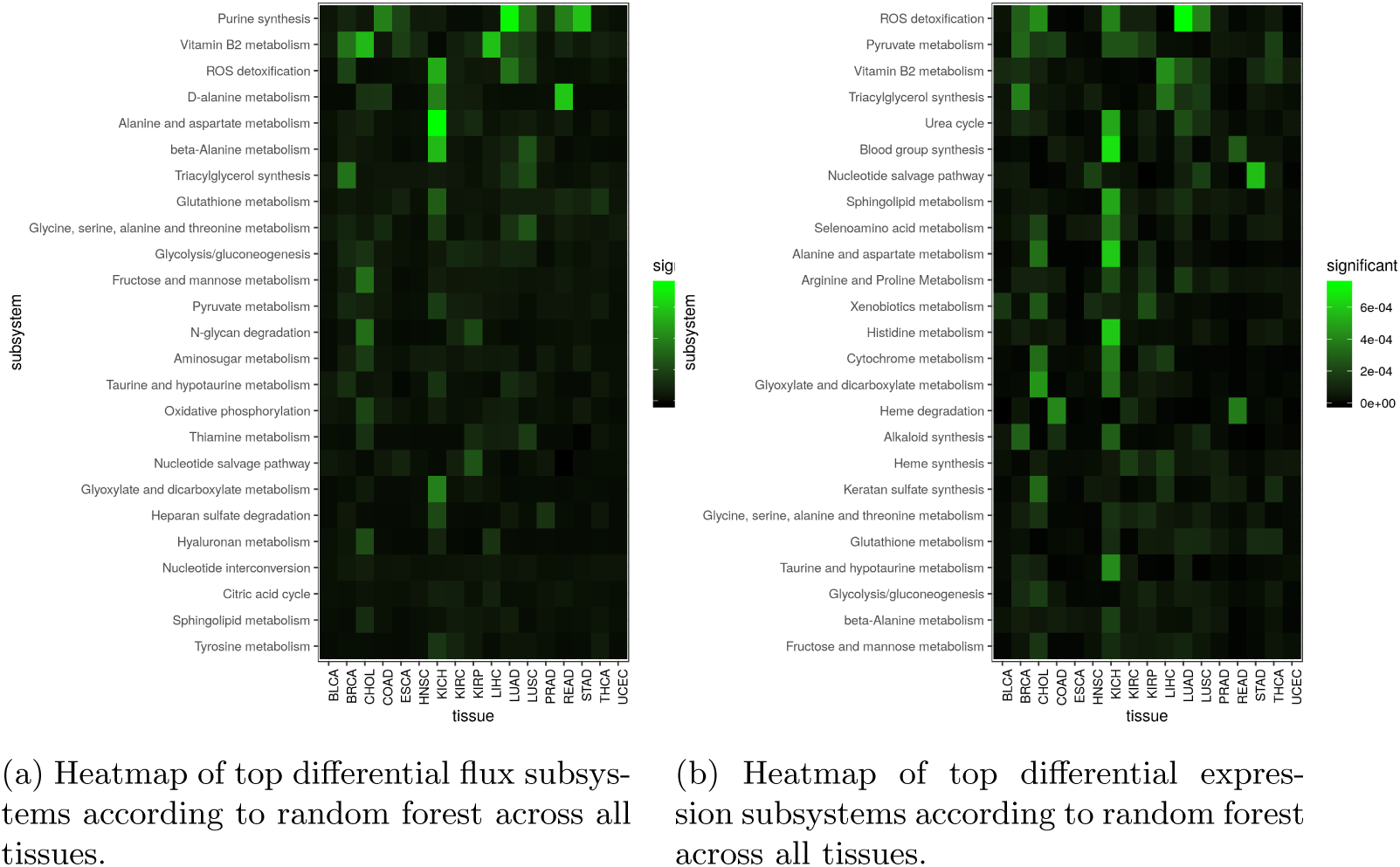
Heatmaps of top differential subsystems by random forests.

**Figure 8:**
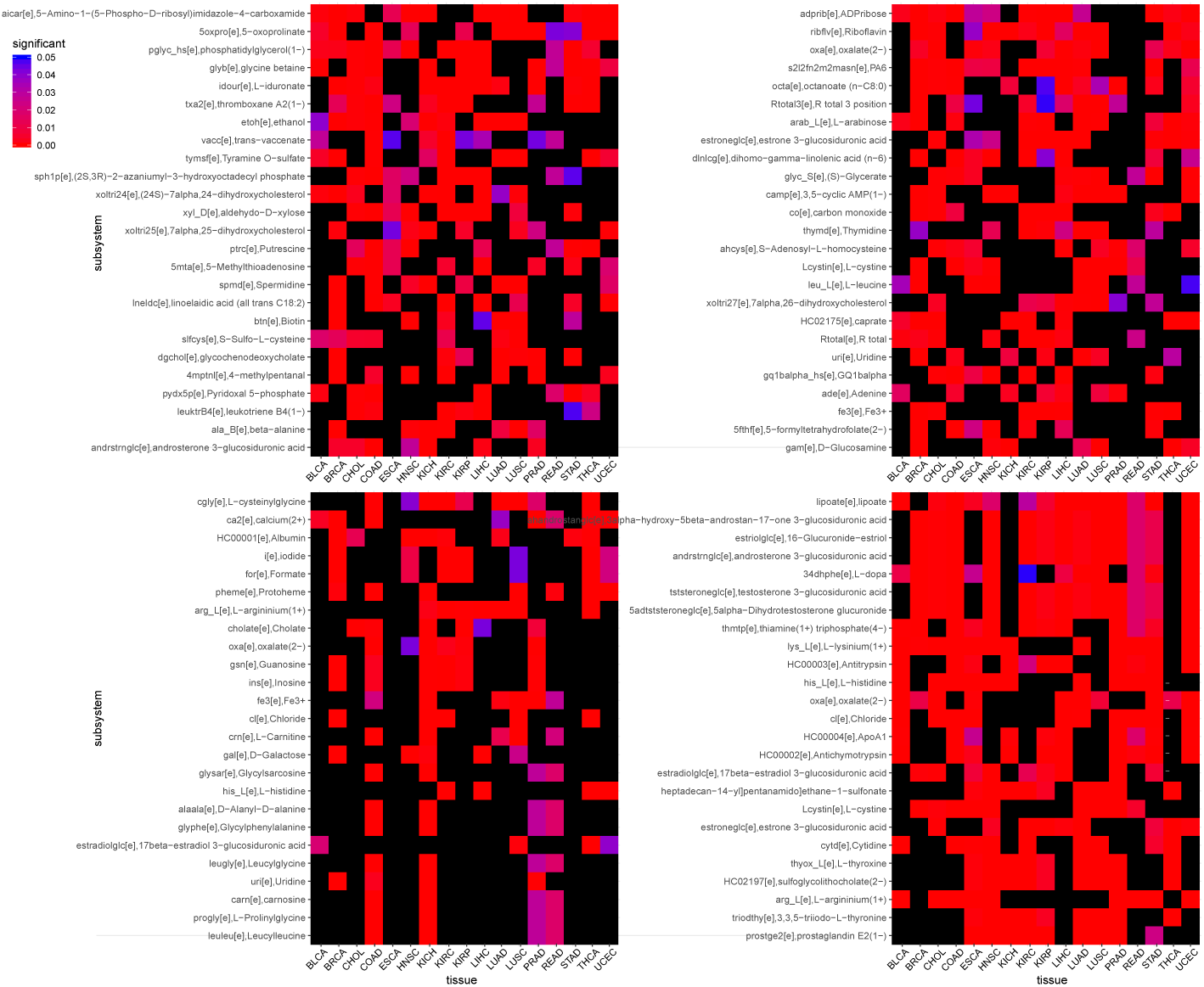
Top-left: Positive differential flux transport reactions. Top-right: Negative differential flux transport reactions. Bottom-left: Positive differential expression reactions. Bottom-right: Negative differential expression reactions.

The most important subsystems for flux differentiation by this metric include a few overlaps with the GSEA results. These include aminosugar metabolism, N-glycan degradation, and fructose and mannose metabolism. However, most of the highest-ranked flux subsystems are different from the GSEA results. They include some that are likely important for tumorigenesis, such as ROS detoxification and glutathione metabolism, which help deal with greatly elevated ROS levels in tumors [2], and vitamin B2 metabolism, which is a critical cofactor for several key metabolic enzymes [30]. Notably, ROS detoxification and B2 metabolism are also important when ranking based on average random forest importance using expression data.

Finally, we also wanted to test for differential flux in transport of extracellular metabolites in tumor vs. normal control tissues. We wanted to focus specifically on these metabolites, because uptake or release of specific metabolites can sometimes be measured in vivo in cancer patients, and may serve as a crucial biomarker. An important example of this is the ingestion of labeled glucose tablets, whose uptake by cancer cells can be monitored by FDG-PET to measure the rate of glucose uptake in general, a key measure of tumor malignancy [6]. Therefore, we examined the top 30 metabolites that most commonly had differential flux or expression, according to the Wilcoxon signed-rank test, across all 17 tissues.

For metabolites whose uptake into cancer cells was predicted to be greater, we observed that among the most common predictions were two biosynthetic compounds, AICAR, an essential intermediary in purine synthesis, and phosphatidylglycerol, a component in the head groups of many membraine lipids. Also common were thromboxane, an eicosanoid, and putrescine, a compound related to spermidine, which may play roles in cancer signaling networks.

Among metabolites whose uptake into cancer cells was predicted to be less, there were two adenine-related compounds, ADP-ribose and cyclic-AMP. cyclic-AMP also plays a major role in many cell signaling pathways. Additionally, arabinose, octanoate, and two compounds related to linolenic also had less flux, suggesting that cancer cells utilize these potential nutrients less.

## 5 Discussion

Altered metabolism is known to be one of the major characteristics of cancer tissues compared to non-cancerous tissues. Despite numerous small-scale experimental studies, changes in metabolic flux in cancer have never been studied experimentally on a genome-wide scale before. Therefore, we applied a computational method called FALCON, developed in our lab previously [12], with the previously published human metabolic model Recon2 [15], to infer flux differences from RNA-seq expression data in the TCGA project.

We first identified several interesting relationships between expression and inferred flux in the TCGA dataset. Although the correlation is in general weak, it is much stronger among the subset of reactions in Recon2 that always have nonzero expression and flux. Furthermore, this correlation is considerably stronger in tumor tissues than their normal control counterparts, and tumor tissues also display a significant smaller number of reactions with nonzero expression and flux. These results imply that tumor tissues have a more streamlined metabolism, with fewer layers of regulation between expression and flux, especially among the most active subset of metabolic reactions.

Our work also helps to identify a large number of candidate metabolic pathways that may be altered in cancer tissues. Although further experimental work must be done to validate whether any of these represent meaningful differences, many of them have previously been shown to involved in tumorigenesis. Furthermore, we also used two different methods for determining differential flux subsystems, GSEA and random forests, and also determined differential expression subsystems. We can use results from all these methods together to focus on subsystems that are important by all measures, and therefore more likely to reflect true biological importance.

Finally, we also tested for significant metabolites whose exchange fluxes are different between normal and tumor tissues. These results also suggest numerous candidate metabolites whose uptake or excretion may be different between cancer and normal tissues.

In total, our work attempts to determine significant metabolic flux differences between cancer and tumor tissues, based on flux inference by the FALCON algorithm. We observed several major pathways that have both differential flux and expression, as well as several metabolites whose uptake and release are predicted to be different, and are linked with the differential pathways. We expect that future studies may further improve the accuracy of metabolic inference in cancer.

